# Autonomic modulations to cardiac dynamics in response to affective touch: Differences between social touch and self-touch

**DOI:** 10.1101/2024.02.20.581146

**Authors:** Diego Candia-Rivera, Rebecca Boehme, Paula C. Salamone

## Abstract

The autonomic nervous system plays a vital role in self-regulation and responding to environmental demands. Autonomic dynamics have been hypothesized to be involved in perceptual awareness and the physiological implementation of the first-person perspective. Based on this idea, we hypothesized that the autonomic activity measured from cardiac dynamics could differentiate between social touch and self-touch. In our study, we used a newly developed method to analyze the temporal dynamics of cardiac sympathetic and parasympathetic activities during an ecologically valid affective touch experiment. We revealed that different types of touch conditions—social-touch, self-touch, and a control object-touch—resulted in a decrease in sympathetic activity. This decrease was more pronounced during social touch, as compared to the other conditions. Following sympathetic decrease, we quantified an increase in parasympathetic activity specifically during social touch, further distinguishing it from self-touch. Importantly, by combining the sympathetic and parasympathetic indices, we successfully differentiated social touch from the other experimental conditions, indicating that social touch exhibited the most substantial changes in cardiac autonomic indices. These findings may have important clinical implications as they provide insights into the neurophysiology of touch, relevant for aberrant affective touch processing in specific psychiatric disorders and for the comprehension of nociceptive touch.

## I. INTRODUCTION

Emotion and stress regulation are key aspects of physiological stability. The intricate physiological mechanisms underlying emotions and their regulation not only affects mental health but also shapes our affective processing and social interactions [1]. Two key mechanisms closely related to this regulation are allostasis and interoception. Allostasis refers to the process of anticipating stressors and achieving stability through physiological adaptations, while interoception refers to the communication within physiological systems, which can be assessed through the body’s internal sensations [2]. The dynamic interplay between allostasis and interoception plays a crucial role in fostering bidirectional communication between our internal sensations and the body’s adaptive responses [3]. In these intricate physiological regulation mechanisms, the role of affective touch emerges as a fundamental factor, as it conveys information from the skin to the central nervous system mediated by the peripheral nervous system [2]. These interoceptive mechanisms may play a significant role in the emotional impact of various forms of affective touch, and ultimately influencing allostatic processes [4].

In a broader terms, the impact of somatosensory stimuli on autonomic dynamics has been superficially explored [5], primarily through the assessment of heart rate variability (HRV). However, this exploration lacks a time-resolved approach which may allow for a comprehensive analysis of the physiological dynamics. HRV serves as a convenient proxy for analyzing autonomic dynamics, given that sympathetic and parasympathetic activations are reflected in alterations in the release of noradrenaline and acetylcholine. These biochemical changes are subsequently reflected in the contraction rate of pacemaker cells [6]. For instance, thermoception can lead to dynamic changes in HRV from an increased sympathetic activity and decreased parasympathetic activity [7], as similarly occurs with nociception [8]. However, the relationship between touch and HRV might be more intricate due to the involvement of an affective component. So far, evidence indicates that affective touch can increase HRV, suggesting a parasympathetic modulation to cardiac dynamics and a sympathetic inhibition [9], [10], whereas sympathetic activations rather occur under the conditions of thermoception and nociception [11], [12]. Pathological conditions can cause disruptions in autonomic balance, such as major depression [13] or depersonalization disorder [14], which could potentially explain the physiological underpinnings of the impairment in the processing of affective touch in those conditions [15], [16].

The distinct contribution of interoceptive inputs to cognition has been hypothesized across various contexts. These contexts encompass regulatory and adaptive processes including fight-or-flight responses or social interactions [17], [18], as well as their potential role in fostering the essential first-person perspective required for perceptual awareness and self-other distinctions [19]. To enhance our understanding of the peripheral dynamics triggered by affective touch, we conducted an experiment involving social and self-produced touch. We aimed to examine the extent to which cardiac sympathetic and parasympathetic activities are differentially modulated under social and self-touch. To our best knowledge, the differences in the autonomic modulations to cardiac dynamics between social and self-touch have not yet been studied. This research gap is significant for comprehending the neurophysiology of affective touch, as the pathways from the skin involve both vagal and sympathetic nerve fibers, which, in turn, can impact cardiac rhythms. In this study, healthy adults underwent an experimental protocol involving social touch, self-touch, and object-mediated touch as a control measure [20]. Given that self-other distinction is believed to entail the integration of tactile, proprioceptive, and interoceptive inputs [21], we hypothesized that cardiac activity could function as a biomarker, facilitating the differentiation between these conditions. Our specific aim was to delineate both cardiac sympathetic and parasympathetic activity, as both may exhibit dynamic activation and deactivation during affective touch [22]. However, the majority of HRV measures lack specificity in identifying their sympathetic or parasympathetic origin. To address this limitation, we employed a recently developed method that offers a robust, time-resolved estimation of cardiac sympathetic and parasympathetic indices from electrocardiograms [23]. This approach enabled us to dynamically quantify the cardiac sympathetic and parasympathetic changes at different latencies with respect to the touch onset, and the differences in these dynamics with respect to the different types of touch.

## II Materials AND Methods

### A. Participants and experimental protocol

A total of 28 healthy adult volunteers participated in this study (16 females, mean age 29.04 years, SD=5.16). Participants were required to be fluent in English, and had no current cardiac, sensory/motor, or affective/psychiatric conditions. Data acquisition was performed at the Center for Social and Affective Neuroscience (CSAN), Linköping, Sweden. All participants provided informed consent in accordance with the Helsinki Declaration and were compensated for their participation. The study was approved by the national ethics board (DNR 2022-02110-01).

The participants engaged in an established experimental task known as the self-other-touch paradigm [20] with modified timing to optimize data for the current research question. The task employed a randomized block design and encompassed three distinct conditions: social touch (being stroked on the left forearm by the experimenter), self-touch (stroking of the participant’s own left forearm), and object-touch (participant stroking a pillow). Each of the three conditions lasted 180 seconds. For the self and object-touch condition, participants were instructed to gently stroke their left arm, mimicking the touch they would use when interacting with someone they like, using their right hand.

Instructions for each block of the task were presented on a screen. The instructions, provided in English, were displayed including the following prompts: “Social touch: Your arm will be touched by the experimenter**»**, «Self-touch: Please stroke your arm”; “Object-touch: Please stroke the object”; Participants either received stimulation or performed the stimulation themselves, continuing for a duration of 180 seconds. The female experimenter (P.C.S), stationed adjacent to the participant replicated the participant’s movements in the same area of the forearm as closely as possible. Participants performed the task with their eyes closed, therefore the experimenter informed them when the condition was over.

The ECG recordings were obtained using *BIOPAC B-Alert* additional 2 channels dedicated to record ECG from the chest.

Recordings were performed with *Acqknowledge* software (*Biopac*) at 2000 Hz.

### B. ECG preprocessing

The ECG data were bandpass filtered with a Butterworth filter of order 4 between 0.5 and 45 Hz. The R-peaks from the ECG were identified using an automatized process, followed by an automated inspection of misdetections and manual correction if required. The procedure was based on Pan-Tompkins algorithm for detecting R-peaks [24]. For the correction of misdetection, all the detected peaks were visually inspected over the original ECG, along with the marks on ectopic heartbeats and the inter-beat intervals histogram.

### C. Computation of cardiac sympathetic and parasympathetic indices

Estimating sympathetic and parasympathetic activity from HRV poses a notable challenge due to the intricate interplay of these autonomic branches within the HRV series. HRV is traditionally studied in the frequency domain, i.e., by identifying and quantifying the amount of slow (<0.15 Hz) and fast (0.15-0.4 Hz) fluctuations in HRV on time [25]. However, the frequency domain approach has been challenged due to the difficulty of associating specific oscillations to either sympathetic or parasympathetic modulations and for not accounting for individual differences in these oscillations [26]. Therefore, untangling these sympathetic and parasympathetic contributions to HRV has prompted the development of alternatives to the frequency domain methods [27], [28].

In this study, a new approach based on Poincaré plot was used to estimate the sympathetic and parasympathetic modulations to HRV [23]. Poincaré plot is a method that depicts the fluctuations on the duration of consecutive inter beat intervals (IBI), accounting for potential nonlinearities in these fluctuations [29], as shown in Fig. 1. The Poincaré plot geometry has been used to characterize changes on HRV by quantifying the ratios of the ellipse formed from consecutive changes in IBIs, representing the short- and long-term fluctuations of HRV, respectively [30]. Importantly, this approach allows to quantify slow and fast fluctuations of HRV at an individual level, without relying on fixed frequency band delimitations. We quantified three distinct features from Poincaré plot: First, the distance of the center of the ellipse to the origin (R), which represents the changes in the baseline heart rate that is increased by sympathetic and decreased by parasympathetic modulations. Second, the minor ratio of the ellipse (SD1), which represents the fast fluctuations of HRV that are triggered by parasympathetic modulations. Third, the major ratio of the ellipse (SD2), which represents the slow fluctuations of HRV that are triggered by sympathetic modulations [23], [30].

**Fig. 1.**
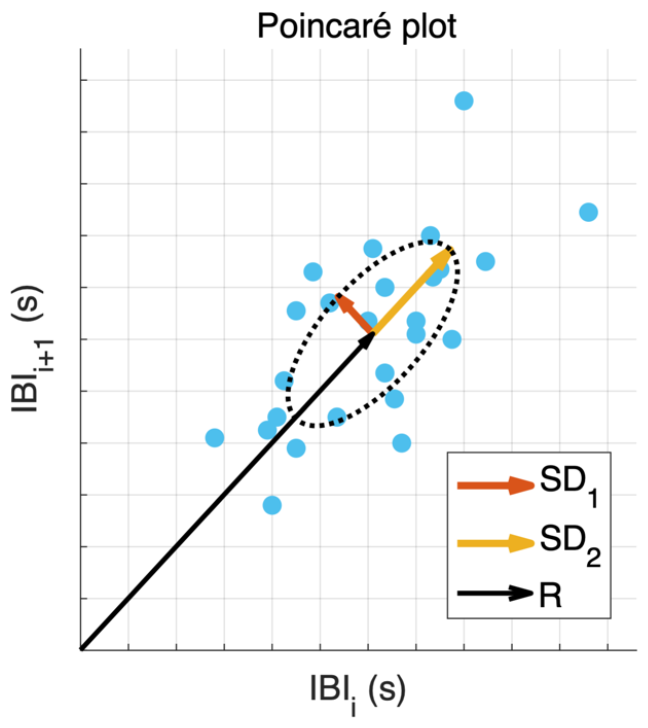
Example of a Poincaré plot. Inter beat intervals (IBIs) are plotted with respect to the next IBI duration. The ellipsoid formed contains three features extracted: the minor ratio (SD1), the major ratio (SD2), and the distance to the origin (R). R represents changes in baseline heart rate that increases by sympathetic and decreases by parasympathetic modulations. SD1 represents the fast fluctuations of HRV that are triggered by parasympathetic modulations. SD2 represents the slow fluctuations of HRV that are triggered by sympathetic modulations.

The time-varying fluctuations of the distance to the origin and the ellipse ratios were computed with a sliding-time window, as shown in Eq. 1, 2 and 3:

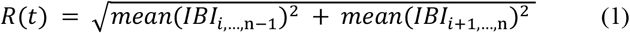

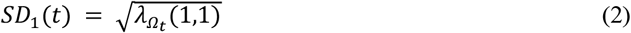

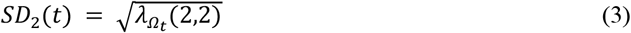

where 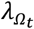 is the matrix with the eigenvalues of the covariance matrix of *IBI*_I,…,n-1_ and *IBI*_i+1,…,n_, with Ω,: *t* – *T* ≤ *t*_i_ ≤ *t*, and *n* is the length of IBI in the time window *Ω*_*t*_.

The distance to the origin *R*_-_ and ellipse ratios *SD*_01_ and *SD*_02_ for the whole experimental duration are computed to re-center the time-resolved estimations of R, SD1 and SD2. Then, the Cardiac Parasympathetic Index (*CPB*) and the Cardiac Sympathetic Index (*CSI*), are computed as follows:

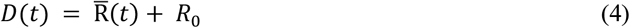

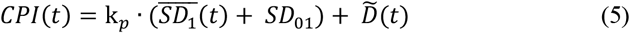

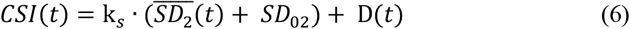

where 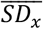 is the demeaned *SD*_*x*_ and 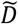 is the flipped *D* with respect the mean. The coefficients k_*p*_ and k_*s*_ define the weight of the fast and slow HRV oscillations, with respect the changes in the baseline heart rate. The values were defined as k_*p*_ = 10 and k_*s*_ = 1, as recommended in the original study [23]. Note that the weights of heart rate and HRV components represent the known effects of autonomic modulations on cardiac dynamics, where sympathetic modulations primarily influence baseline heart rate [31], but also slow HRV changes [32], and parasympathetic modulations are mainly associated to fast HRV changes [32].

For a comprehensive description of the model, see [23]. The method was previously validated in different standard conditions where cardiac sympathetic and parasympathetic activities are modulated, including postural changes, cold-pressor test, and physical exercise test. The software can be gathered from *https://github.com/diegocandiar/robust_hrv*.

### D. Statistical analysis

Within-subject statistical analyses between the three touch conditions were performed through nonparametric Wilcoxon signed-rank tests. The statistical testing was performed per autonomic index, in which the inputs correspond to the averaged CSI or CPI in a defined interval during the different experimental conditions. The significance level of p-values was corrected in accordance with the Bonferroni rule for the three possible paired comparisons, with an uncorrected statistical significance set to α = 0.05. The samples were described group-wise using the medians ± median absolute deviation (MAD), where MAD (X) = Median (|X − Median (X)|). The time-resolved autonomic indices were z-score normalized on time for the whole experimental recording, prior segmentation for statistical analyses.

Additionally, a multivariate analysis was included for ranking the contributions of cardiac autonomic dynamics at different latencies for the distinction of the different types of touch included in this study. The ranking computation was based on Minimum Redundancy Maximum Relevance (MRMR) scores [33], in which a higher score indicates a higher relevance indistinguishing between classes.

## III. Results

We examined cardiac sympathetic and parasympathetic indices obtained from ECG recordings in a study involving healthy individuals undergoing an affective touch protocol. The protocol included social touch, self-touch, and object-touch. The computation of cardiac sympathetic and parasympathetic indices was performed using a robust method based on fluctuating Poincaré plots, which has been demonstrated to be a compelling approach to accurately perform these estimations [23]. The time-resolved estimation of autonomic indices for each condition is presented in Fig. 2. In all three conditions, a decrease in cardiac sympathetic indices was observed, while there was a shorter increase in cardiac parasympathetic activity at an early stage of the touch (during the first 20 seconds).

**Fig. 2.**
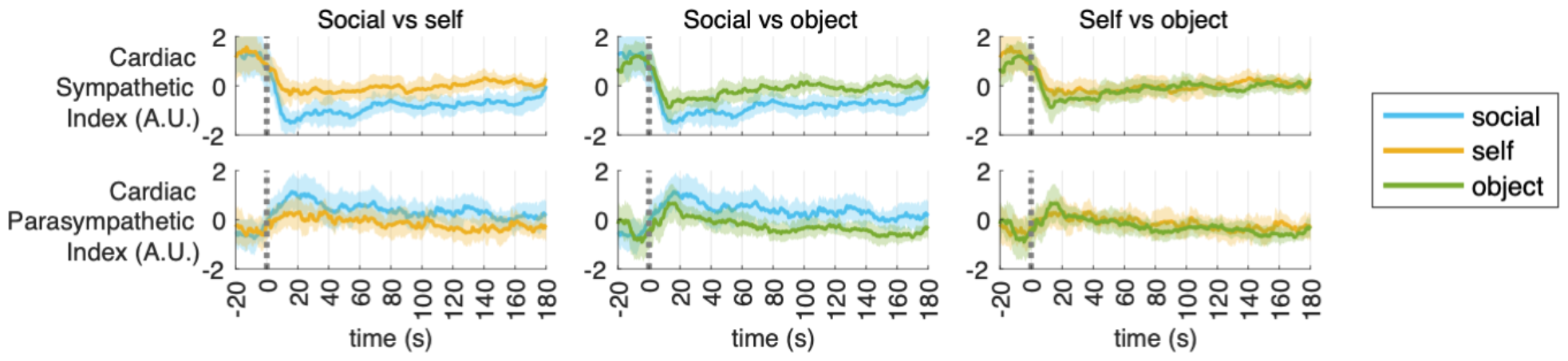
Time course of the changes in cardiac sympathetic (CSI) and parasympathetic indices (CPI) in response to social-touch, self-touch, object-touch. Time course represents the group median and shaded areas the median absolute deviation (MAD) of CSI and CPI.

We conducted a comparison of the three touch conditions at three specific intervals: 0-60s, 60-120s, and 120-180s, to further describe the autonomic dynamics that allow the affective touch distinctions on time. Significant differences in sympathetic indices were observed in all three intervals when comparing social touch to self-touch and social touch to object touch (Fig. 3, Table I). However, there were no significant differences at any interval when comparing self-touch to object touch.

**Fig. 3.**
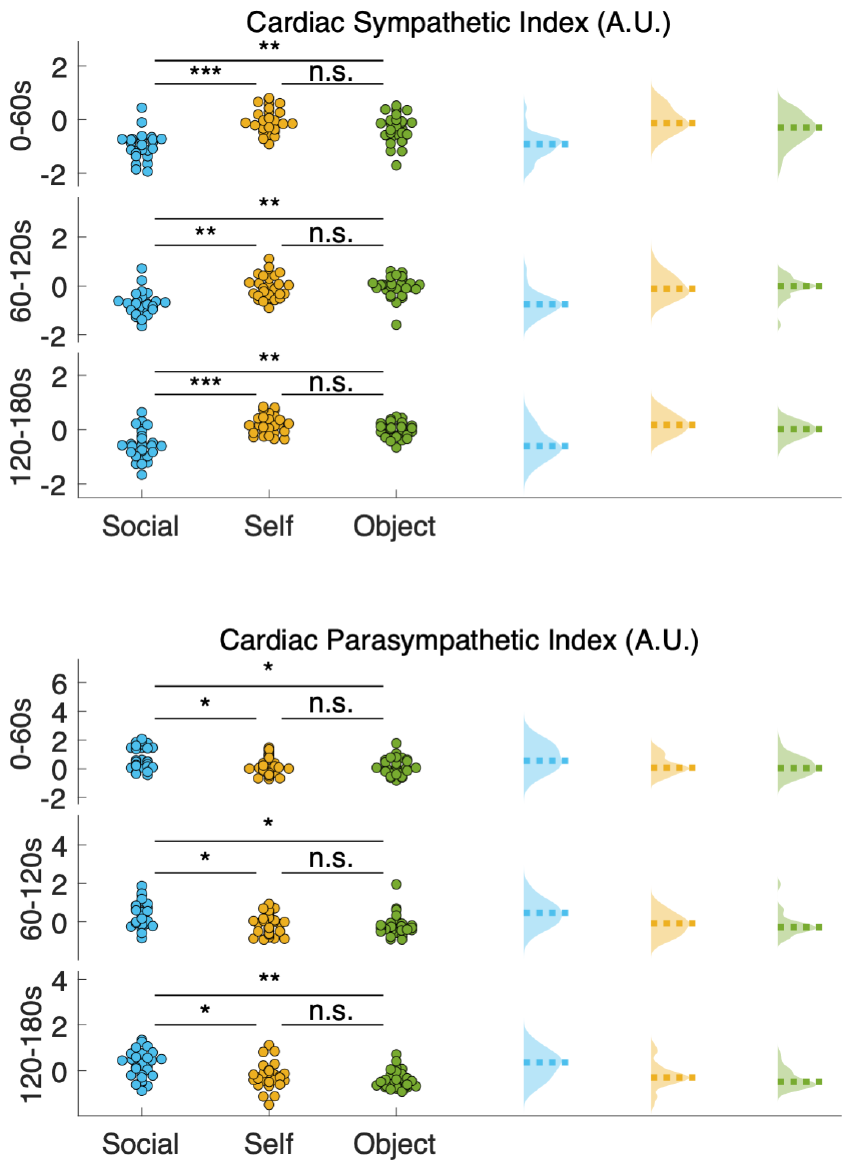
Paired comparisons of cardiac sympathetic indices between social-touch, self-touch, object-touch. Each data point corresponds to one participant at the defined time interval. The comparisons were performed on 1-minute averages in the intervals 0-60, 60-120 and 120-180s. P-values correspond to the Wilcoxon signed-rank test. Significance is defined by the Bonferroni rule at =0.0166. A.U.: Arbitrary Units; n.s.: non-significant

**TABLE 1.** Paired tests between self, social, and object touch.

| Cardiac Sympathetic Index |  |  |  |
| --- | --- | --- | --- |
|  | Self vs Social | Social vs Object | Self vs Object |
| 0 – 60 s | Z=4.3493,<br>p<0.0001 | Z=3.3929,<br>p=0.0007 | Z=1.7762,<br>p=0.0757 |
| 60 – 120 s | Z=3.6434,<br>p=0.0003 | Z=3.7117,<br>p=0.0002 | Z=0.2277,<br>p=0.8199 |
| 120 – 180 s | Z=4.0305,<br>p<0.0001 | Z=3.7117,<br>p=0.0002 | Z=1.8673,<br>p=0.0619 |

| Cardiac Parasympathetic Index |  |  |  |
| --- | --- | --- | --- |
|  | Self vs Social | Social vs Object | Self vs Object |
| 0 – 60 s | Z=2.6415,<br>p=0.0083 | Z=2.5504,<br>p=0.0108 | Z=0.0911,<br>p=0.9274 |
| 60 – 120 s | Z=2.6187,<br>p=0.0088 | Z=3.1425,<br>p=0.0017 | Z=0.6604,<br>p=0.5090 |
| 120 – 180 s | Z=2.4821,<br>p=0.0131 | Z=3.3474,<br>p=0.0008 | Z=1.3663,<br>p=0.1718 |

Paired Wilcoxon tests were performed on the average of cardiac sympathetic indices within 0-60s, 60-120s, and 120-180s intervals, with respect to the touch onset. Significance is shown in shaded gray areas and it was defined by the Bonferroni rule at $\alpha=0.0166$ .

Similar to the sympathetic indices, we found significant changes in the cardiac parasympathetic indices across all three intervals when comparing social touch to self-touch and social touch to object touch. However, there were no significant differences at any interval when comparing self-touch to object touch (see Fig. 3, Table I). We found that cardiac sympathetic activity was consistently reduced under social touch, whereas cardiac parasympathetic activity tended to be higher in that condition. Notably, the differentiation between social touch and object touch became more pronounced in the later stages of the protocol.

Expanding upon these findings, we aimed to assess the distinguishability of the experimental conditions in a multivariate space. We conducted a feature ranking analysis using the MRMR algorithm to determine which sympathetic or parasympathetic indices, and at what latency, were more effective in distinguishing the various types of touch encompassed within this study (Table II). The sympathetic (intervals 0-60s and 60-120s) and the later parasympathetic (120-180s interval) modulations appeared among the three most relevant features in distinguishing between all possible paired comparisons. Based on the feature ranking for the distinctions of the three types of touch, we focused on the averaged sympathetic and parasympathetic indices within the aforementioned intervals. As depicted in Fig. 4, cardiac sympathetic and parasympathetic indices provide complementary information to distinguish between the touch conditions, with a more robust separability between social touch and both self and other touch. In this sense, social touch provides a greater modulation of cardiac parasympathetic activity while at the same time showing a greater reduction in cardiac sympathetic activity.

**TABLE 2.** Feature ranking based on minimum redundancy maximum relevance (MRMR) score.

| Feature ranking | Feature (MRMR score) |  |  |
| --- | --- | --- | --- |
|  | Social vs self | Self vs object | Social vs object |
| 1 | Sympathetic 0-60s (0.3526) | Sympathetic 0-60s (2.9e-15) | Sympathetic 60-120s (0.2239) |
| 2 | Parasympathetic 120-180s (0.1920) | Sympathetic 60-120s (2.4e-15) | Parasympathetic 120-180s (0.1574) |
| 3 | Sympathetic 60-120s (0.1920) | Parasympathetic 120-180s (2.4e-15) | Sympathetic 0-60s (4.8e-15) |
| 4 | Sympathetic 120-180s (0.1893) | Parasympathetic 0-60s (2.4e-15) | Sympathetic 120-180s (4e-15) |
| 5 | Parasympathetic 0-60s (3.9e-15) | Parasympathetic 60-120s (1.6e-28) | Parasympathetic 60-120s (2.6e-15) |
| 6 | Parasympathetic 60-120s (2.6e-15) | Sympathetic 120-180s (1.2e-28) | Parasympathetic 0-60s (6.9e-29) |

The scores were computed for all possible comparisons between social, self, and object touch using as features the averages of cardiac sympathetic and parasympathetic indices within 0-60s, 60-120s, and 120-180s intervals.

**Fig. 4.**
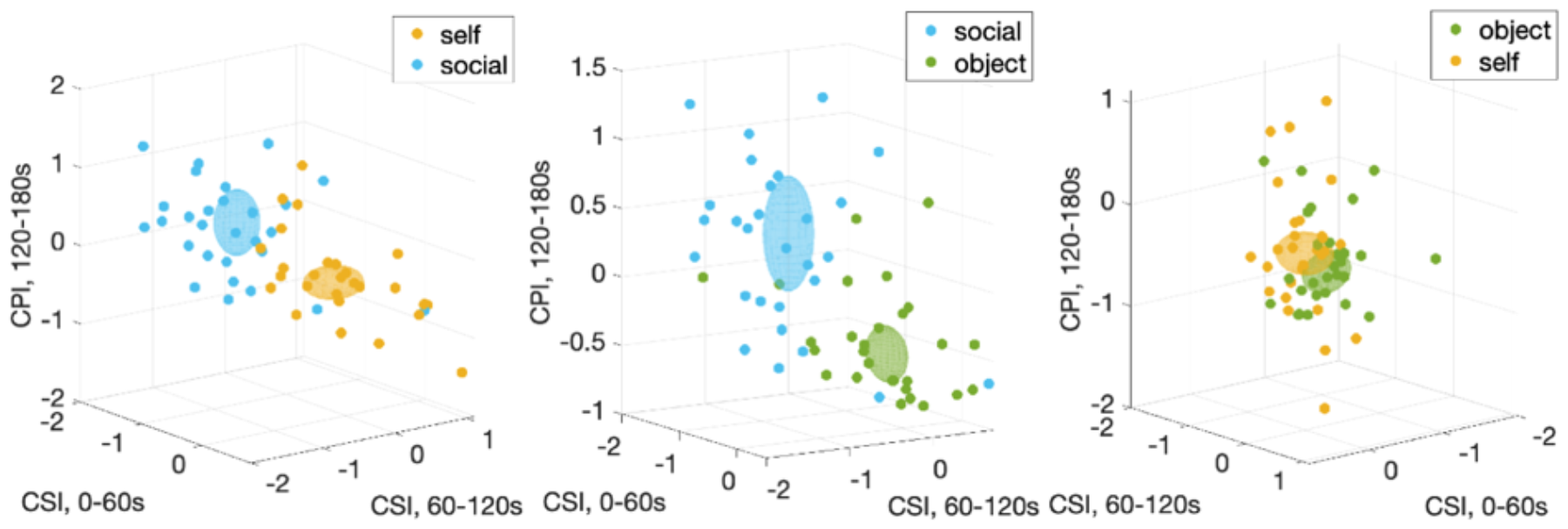
Combined indices for cardiac sympathetic indices (CSI) and cardiac parasympathetic indices (CPI) in 3D scatter plots to distinguish self-touch, social-touch, object-touch. The CSI was considered in the intervals 0-60s and 60-120s, and CPI in the interval 120-180s, with respect to the touch onset. Those intervals represent the instances in which a major separability between the conditions occurred, as quantified in a multivariate analysis. The displayed ellipsoids are centered in the group median and their ratios correspond to the median absolute deviations per dimension.

## IV. Discussion

Understanding the physiological mechanisms underlying the processing of social touch holds significant relevance for various domains, including clinical conditions in which affective touch appears altered. This study marks the pioneering effort to uncover the temporal intricacies of sympathetic and parasympathetic influences on cardiac dynamics across various touch modalities. We demonstrate how social and self-touch elicit distinct autonomic modulations, potentially serving as valuable biomarkers of interest, especially within the field of affective computing. In our study, we utilized a recently developed method [23] to analyze the temporal dynamics of cardiac sympathetic and parasympathetic activities under an affective touch paradigm. Our results revealed that affective touch — social touch and self-touch — elicited a decrease in sympathetic activity. The sympathetic decrease was notably more pronounced during social touch as compared to the other conditions. Moreover, we observed an increase in parasympathetic activity specifically during social touch, which furthered the differentiation from self-touch. Interestingly, as the protocol progressed, the distinction between parasympathetic responses to social touch and object touch became more prominent. Remarkably, by combining the sympathetic and parasympathetic indices, we were able to further differentiate the experimental conditions, with social touch exhibiting the most substantial modulations on cardiac autonomic dynamics.

Sympathetic activity might play a role in facilitating the functioning of tactile receptors, as suggested by findings that the inhibition of sympathetic activity enhances tactile sensitivity [34]. Our results are in line with this effect, which is thought to be triggered by the activation of C-fibers, specialized receptors that respond to slow and gentle touch [9]. In comparison, the faster A-delta fibers in the skin are thought to be responsible for sympathetic activation during thermal and nociceptive stimuli [11]. Additional evidence indicates that sympathetic reactivity, as measured by changes in pupil diameter and skin conductance, is influenced by the velocity of stroking rather than touch itself [35], [36]. The observed effects on sympathetic indices may arise from ascending signaling from the skin to the brain via spinal cord pathways. Animal models have demonstrated that inhibitory influences on sympathetic activity involve pathways ascending through the dorsolateral funiculus and sulcus areas of the spinal cord [37]. It is noteworthy that processing in the spinal cord also contributes to the modulation of the sense of body ownership [38] and the distinction between self-touch and social touch [20].

Our present and previous experimental evidence suggests that parasympathetic activations represent the affective component in touch. Cardiac deacceleration modulated from parasympathetic inputs has been reported under affiliative contexts [39]–[41] and in infants [42]. Furthermore, the stimulation of C-fibers within an optimal stroking velocity range produced an increased parasympathetic modulation to cardiac dynamics, which was sustained into the subsequent post-touch period [43]. The observed parasympathetic activity in our result indicates that social touch triggers additional physiological adjustments and involves active integration of interoceptive inputs. Previous studies have linked variations in parasympathetic tone to fluctuations in attention and emotional processing [44]. In fact, the differences in parasympathetic modulations to cardiac dynamics under emotion regulation tasks have demonstrated its ability to distinguish healthy individuals from those with certain pathologies [45], [46].

The sympathetic and parasympathetic dynamics have been previously associated with behavior in the polyvagal theory [17], which describes neural circuits involved in homeostatic regulation and adaptation. According to the polyvagal theory [17], the parasympathetic branch is primarily associated with promoting social engagement, as it correlates with reactivity, emotional expression, and self-regulation skills. Notably, parasympathetic activity has been found to promote emotional processing in response to audiovisual stimuli [47], which supports that subjective emotional experiences may require the ongoing integration of interoceptive inputs in the brain.

Previous research has established connections between brain-heart mechanisms and various cognitive processes such as body ownership, perspective-taking, and consciousness [19], [48]–[52]. Given these connections, it is conceivable that autonomic dynamics play a role in the neural mechanisms that differentiate between social touch and self-produced touch in the brain. In the framework of predictive coding, the integration of interoceptive inputs becomes essential for regulatory and anticipation processes [1], [53]. Somatosensory detection and tactile actions are closely linked to the muscle contraction phase of the cardiac cycle [54]–[56], and neural responses to heartbeats are also related to somatosensory detection [57], [58]. This suggests that interoceptive inputs are integrated during conscious somatosensory perception. Further supporting this idea, the insula — a brain region involved in interoceptive processing—plays a role in distinguishing between observing others’ somatosensory experiences and one’s own somatosensory experiences [59], and its damage disrupts affective touch perception [60]. Finally, the reported relationship of brain-heart interactions with self-other distinctions [61]–[64] and emotion processing [47], [64]–[68] further supports the role of autonomic dynamics in the processing of affective stimuli.

Through a newly developed method, we analyzed the temporal dynamics of cardiac sympathetic and parasympathetic activities during an ecologically valid affective touch experiment. Our findings have important clinical implications, specifically for the treatment of disrupted affective touch perception/processing seen in many psychiatric disorders [69]. Understanding the mechanisms underlying touch pathways is crucial, as disruptions in these pathways can lead to altered sensitivity and specificity of tactile receptors, resulting in the perception of tactile stimuli as uncomfortable or painful [34]. Enhancing our knowledge of the physiological aspects of social touch may offer insights for the treatment of pathological painful touch [70] and make further links with the neural mechanisms of nociceptors [71], [72]. By elucidating the relationship between autonomic activity and touch in the time domain, we can potentially uncover strategies for alleviating pain through touch-based interventions [73]–[75]. Overall, our findings have clinical relevance by highlighting the potential implications for the treatment of disrupted affective touch perceptions and processing.

## V. Conclusion

We revealed that social touch causes a greater decrease in sympathetic activity as compared to other types of touch. Subsequently, an increase in parasympathetic activity during social touch further distinguished its autonomic dynamics. We showed that the combination of sympathetic and parasympathetic indices serves to enhance the recognition of social touch. Our findings may have important clinical implications as they provide insights into the neurophysiological dynamics of touch, which could be relevant for investigating pathological conditions with disrupted affective touch processing as a comorbidity.

